# GALILEO Generatively Expands Chemical Space and Achieves One-Shot Identification of a Library of Novel, Specific, Next Generation Broad-Spectrum Antiviral Compounds at High Hit Rates

**DOI:** 10.1101/2025.01.17.633620

**Authors:** Tyler Umansky, Virgil Woods, Sean M. Russell, David S. Garvey, Davey M. Smith, Daniel Haders

## Abstract

The COVID-19 pandemic (2019-2023) demonstrated the need for safe and effective, stockpiled broad-spectrum antiviral drugs to suppress unexpected viral outbreaks. The inability of the pharmaceutical industry to create such a therapeutic in the 6 years since the onset of COVID-19 demonstrates antiviral drug development must undergo a paradigm shift for this to occur. AI-based target and medicinal chemistry discovery platforms such as GALILEO and its geometric graph convolutional network tool ChemPrint, which we recently published, hold promise in accelerating and reimagining the drug development process. GALILEO identified the Thumb-1 site (an allosteric subdomain of the viral RNA polymerase) to be structurally conserved across numerous viral species and MDL-001 (an orally available therapeutic with a favorable pharmacokinetics and safety profile in humans) to be a potent inhibitor thereof. Published preclinical proof-of-concept studies demonstrated MDL-001 as a first-in-class broad-spectrum antiviral drug. This study leverages GALILEO’s generative and multimodal discovery tools to create trillions of new chemical entities (NCEs) from MDL-001’s pharmacophoric scaffold and select a library of highly specific and optimized compounds for next-generation broad-spectrum antiviral development. Specifically, ChemPrint’s one-shot predictions identified 12 NCEs with predicted affinity to Thumb-1 and significantly reduced, or no, affinity to MDL-001’s original target. *In vitro* bioassays demonstrated a 100% hit rate, with all 12 NCEs having antiviral activity against Hepatitis C Virus (HCV) and/or human Coronavirus 229E. *In vitro* studies also demonstrated reduced activity of 800-fold to greater than 15,000-fold relative to MDL-001’s originally designed mechanism of action (MoA). In Tanimoto similarity plots, the 12 NCEs lacked chemical relatedness to known antiviral drugs, including MDL-001 (average Tanimoto coefficient: 0.38) and Beclabuvir, (average Tanimoto coefficient: 0.13) a HCV Thumb-1 ligand that lacks broad-spectrum activity. This study showcases GALILEO’s ability to generate vast NCE libraries and ChemPrint’s extrapolative capabilities to discover large, potent NCE libraries of compounds, specific to a complex target that are novel to known chemistry at high hit-rates.

## Introduction

From the Justinian plague in the premodern times (500s A.D.), to the COVID-19 outbreak in 2019, the history of humankind can be charted along the timeline of pandemic outbreaks.^1^ Zoonotic viruses, particularly RNA viruses, or viruses with an RNA step, remain the primary pandemic catalyzers of the modern era (e.g., Ebola virus; human immunodeficiency virus; multiple influenza A virus subtypes, and Coronavirus variants).^1,2^ In the late 1890s, an undefined influenza A virus subtype triggered the first modern pandemic of the modern era (the so-called Russian pandemic).^2^ Multiple influenza A virus subtypes followed suit afterward, triggering several pandemics throughout the 1900s and early 2000s.^3^ The Great flu in 1918 (influenza A/H1N1 virus), brought worldwide death in epic proportions, only matched by the COVID-19 pandemic in 2019.^1^ Just shy of 25 years into the 21^st^ century, Coronaviruses have made a mark with two pandemics: the severe accurate respiratory syndrome (SARS) pandemic (2002-2004), and the COVID-19 pandemic (2019-2023), mentioned above.^1^

In the aftermath of the COVID-19 pandemic, epidemiological surveillance programs have shown that zoonotic virus emergence is far from winding down.^4^ Vaccines have been determinant in keeping contemporary influenza outbreaks at bay in many, but not all, cases and more recently, managing the COVID-19 pandemic.^5,6^ Vaccine development is, however, a lengthy and often impossible task, usually initiated only after proof of a virus’s pandemic potential.^6^ Plus, vaccine storage and distribution can be an obstacle.^6^ Small-molecule antiviral drugs, the alternative therapeutic approach, often have a limited activity spectrum.^7^ Before COVID-19 vaccines were developed, available small-molecule antivirals did little to abate the COVID-19 pandemic’s heavy death toll.^8^ This grim experience underlined the need to develop safe and effective, stockpiled, broad-spectrum antiviral drugs that can be implemented at the very onset of a viral outbreak.

We set out to change the antiviral drug discovery paradigm in two significant ways: 1) by cutting the time and cost to reach the clinical stage and 2) by moving away from the one virus, one drug approach, which has inhibited broad-spectrum antiviral development.^9^ We recently developed GALILEO, an artificial intelligence (AI) drug discovery suite featuring ChemPrint⎯a graph convolutional network, which learns from geometric graph embeddings^10,11,12^. In a previous study, leveraging GALILEO’s bioinformatics tools and ChemPrint, we found that MDL-001 (originally a selective estrogen receptor modulator with demonstrated safety in rodents and humans at high concentrations for extended durations) targets the Thumb-1 pocket in the RNA-dependent RNA polymerase (RdRp) encoded by Hepatitis C Virus (HCV)^11,12^. We also found that the Thumb-1 pocket is structurally conserved in many pathogenic viruses, making it a druggable target for broad-spectrum antiviral compounds, such as MDL-001^11,12^.

In the present study, we aimed to build on the success of MDL-001, a state-of-the-art broad-spectrum antiviral compound with exceptional efficacy and safety, by leveraging our GALILEO platform to create a library of next generation, antiviral NCEs with increased specificity at high hit rates. Using GALILEO’s generative capabilities, we generated 52 trillion NCEs across a large chemical space, starting from MDL-001’s pharmacophoric scaffold. Through a multimodal approach, ChemPrint’s one-shot predictions identified 12 NCEs from GALILEO’s trillion-molecule chemical space that retained MDL-001’s antiviral activity and lacked binding to MDL-001’s original target. When validated *in vitro*, all 12 NCEs exhibited antiviral activity (100% hit rate). Additionally, *in vitro* studies demonstrated a significant increase in target specificity—ranging from 800-fold to greater than 15,000-fold—against MDL-001’s original target. Chemical space maps and Tanimoto coefficients revealed that these 12 NCEs are significantly dissimilar from MDL-001 (average Tanimoto coefficient: 0.38) and a large collection of known antiviral inhibitors, including Beclabuvir (average Tanimoto coefficient: 0.13), a Thumb-1 HCV RdRp inhibitor that lacks evidence of broad-spectrum antiviral activity^12^. This study highlights the power of our one-shot, multimodal approach to systematically discover large, optimized NCE libraries containing potent compounds for complex targets with off-target interactions eliminated at high hit-rates. The study also represents a significant milestone in a multi-generational therapeutic program that began with MDL-001 and continues to drive advancements in NCE discovery and optimization.

These findings go beyond showing that GALILEO represents a new way of discovering drug leads. The workflow presented here underlines the advantages of generative tools like GALILEO, which by creating large chemical spaces, open the door to on-demand drug development programs with a high hit rate and designed target specificity. ChemPrint also brings great value, enabling the one-shot identification of drugs, such as the 12 NCEs we report here, that are optimized for multiple properties, through a multimodal approach.

## Results

The goal of this study was to 1) use GALILEO’s generative tools to create a trillion compound NCE space inspired by the scaffold of the potential first-in-class therapeutic MDL-001 and 2) use ChemPrint’s discovery tools in a one-shot, multimodal approach to identify a next generation library of potent NCEs with antiviral activity that selectively targets the broad-spectrum Thumb-1 pocket in viral RdRp and eliminates off-target activity. We achieved this by combining the computational AI tools embedded in GALILEO with downstream *in vitro* assays (experimental diagram depicted in **Figure 1**).

**Figure 1.**
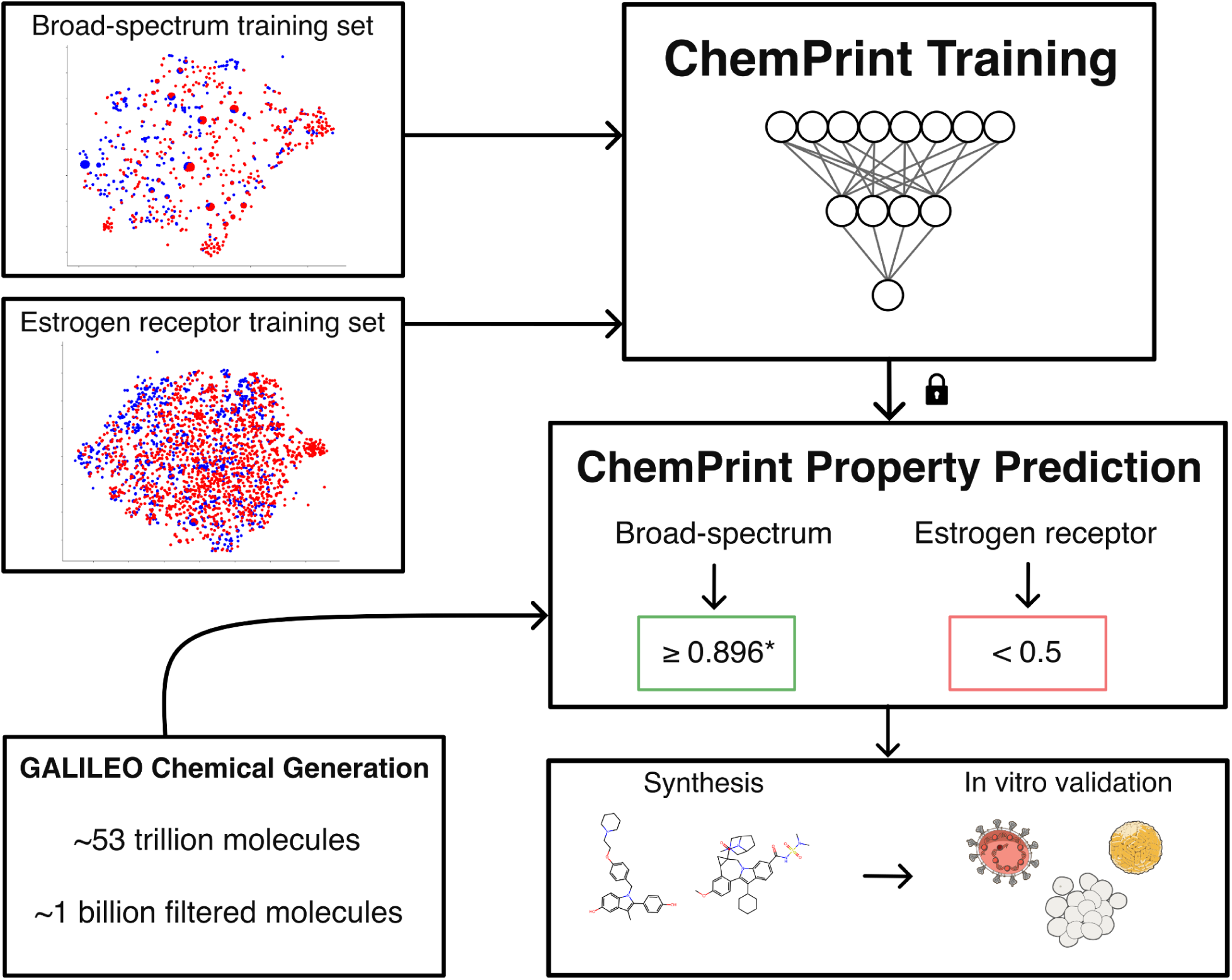
Experimental workflow demonstrating our generative and multimodal drug discovery approach to remove off-target interactions while optimizing antiviral properties.

### Generative chemical space

We leveraged GALILEO to generate a combinatorial chemical space of 52 trillion probable NCE antiviral compounds, by adding a combinatorial set of R group substituents to pharmacophoric scaffolds, inspired from MDL-001’s chemical structure. We next collapsed the combinatorial chemical space by reducing R substituent chemical complexity, resulting in an inference library of 1 billion compounds (see Materials and Methods). The inference library containing 1 billion molecules retained NCE diversity features unique to the combinatorial chemical space; and made the downstream computational tasks more manageable.

We processed the inference library using two ChemPrint models, trained to identify NCEs with a Thumb-1 binding probability higher than that of MDL-001 (0.896); and to remove unspecific estrogen receptor binders, MDL-001’s originally developed MoA (**Figure 1**). ChemPrint is a convolutional neural network that extracts features from small molecules embedded as geometrical graphs. Thus, we embedded the NCEs in the inference library as geometrical graphs, and once embedded, we loaded the inference library dataset to ChemPrint. In the first case, we used the ChemPrint model trained to predict NCE/Thumb-1 interactions (Materials and Methods). The results from this analysis revealed that from the 1-billion inference library input, four million NCEs had a Thumb-1 binding probability ≥ 0.896. We narrowed this list down to 500 structurally diverse NCEs, after removing compounds with challenging synthetic pathways and halogen-reactive moieties.

MDL-001 (the pharmacophoric scaffold precursor of the combinatorial chemical space) is a known modulator of the estrogen receptor^14,15^, raising the possibility that the NCEs may also modulate the estrogen receptor. Following the same approach used to develop ChemPrint for identifying NCEs with the desired property of broad-spectrum Thumb-1 interaction, we trained the model to exclude compounds with undesired properties, specifically estrogen receptor interaction (Materials and Methods). An estrogen receptor binding probability < 0.5 was used to refine the 500-NCE list to three compounds.

Using GALILEO, we generated a second inference library based on the three selected NCEs. This library included synthetic pathway intermediaries, compound isomers, and chemical analogues derived from each NCE. For this second round of ChemPrint filtering, a less stringent probability cutoff of 0.5 was applied to the ChemPrint model predicting Thumb-1 interactors, compared to the 0.896 cutoff used for the initial library. This adjustment was made to prioritize accessibility, allowing for the inclusion of compounds that could be readily synthesized and collected during the development process. After processing this library with the same two ChemPrint models, we obtained a final shortlist of 12 NCEs (**Table 1**).

**Table 1.**
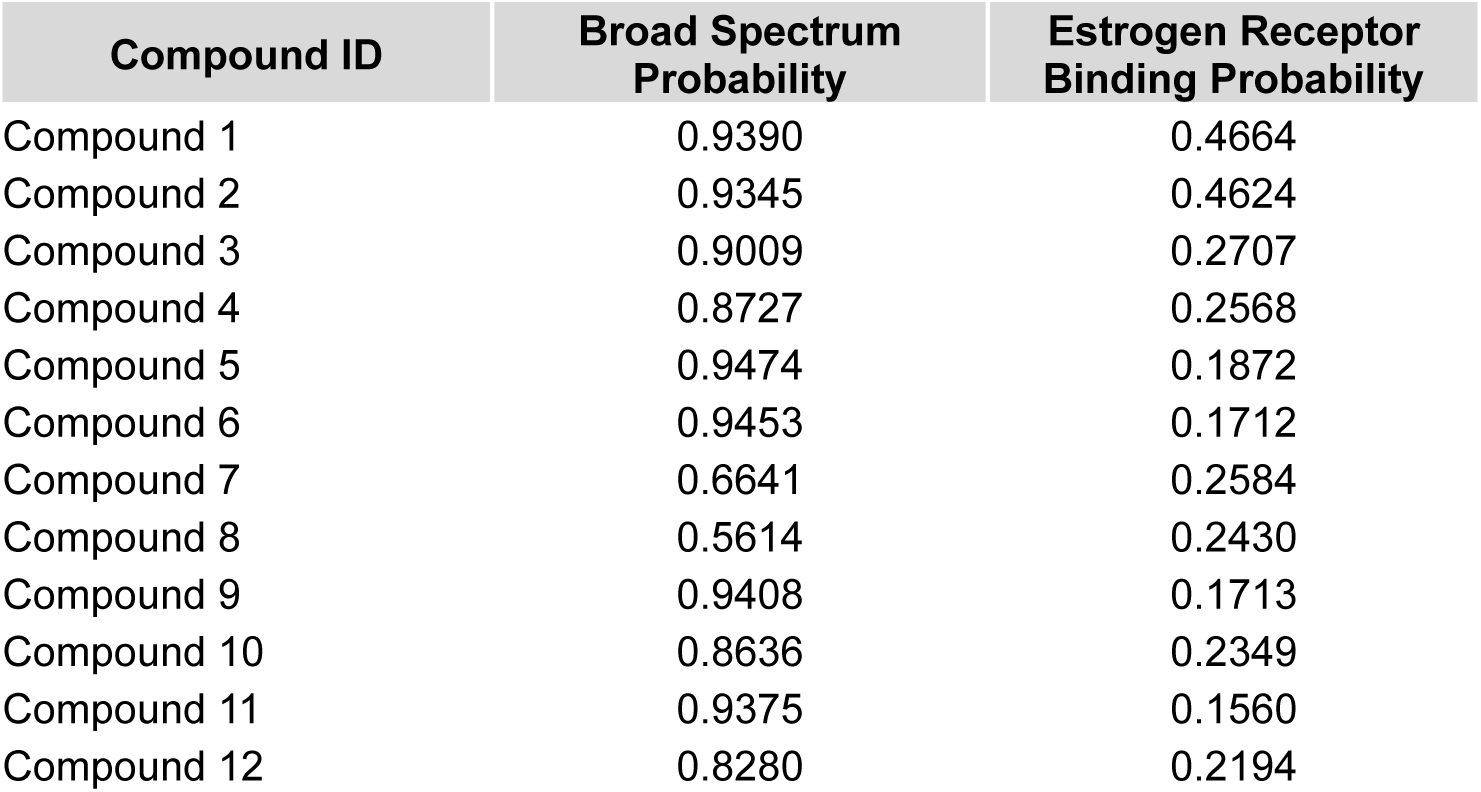
Broad-Spectrum and Estrogen Receptor Binding ChemPrint model predictions of 12 compounds nominated for synthesis.

### Chemical Space Analysis

To evaluate the mechanisms of action (MoA), novelty, diversity, and similarity of the nominated NCEs, we analyzed their chemical space. Chemical space mapping was performed using principal component analysis (PCA)^16^ and t-distributed stochastic neighborhood embedding (t-SNE)^17^ to visualize relationships and Tanimoto coefficients to quantify them. The analysis included MDL-001, Beclabuvir, the 12 NCEs, a subset of ChemPrint-predicted broad-spectrum Thumb-1 compounds, and curated antiviral datasets with known viral RdRp MoAs (labels explained in Material and Methods) (**Figure 2A-B**). Both PCA and t-SNE used only embedded chemical structures as input features, without incorporating MoA labels.

**Figure 2.**
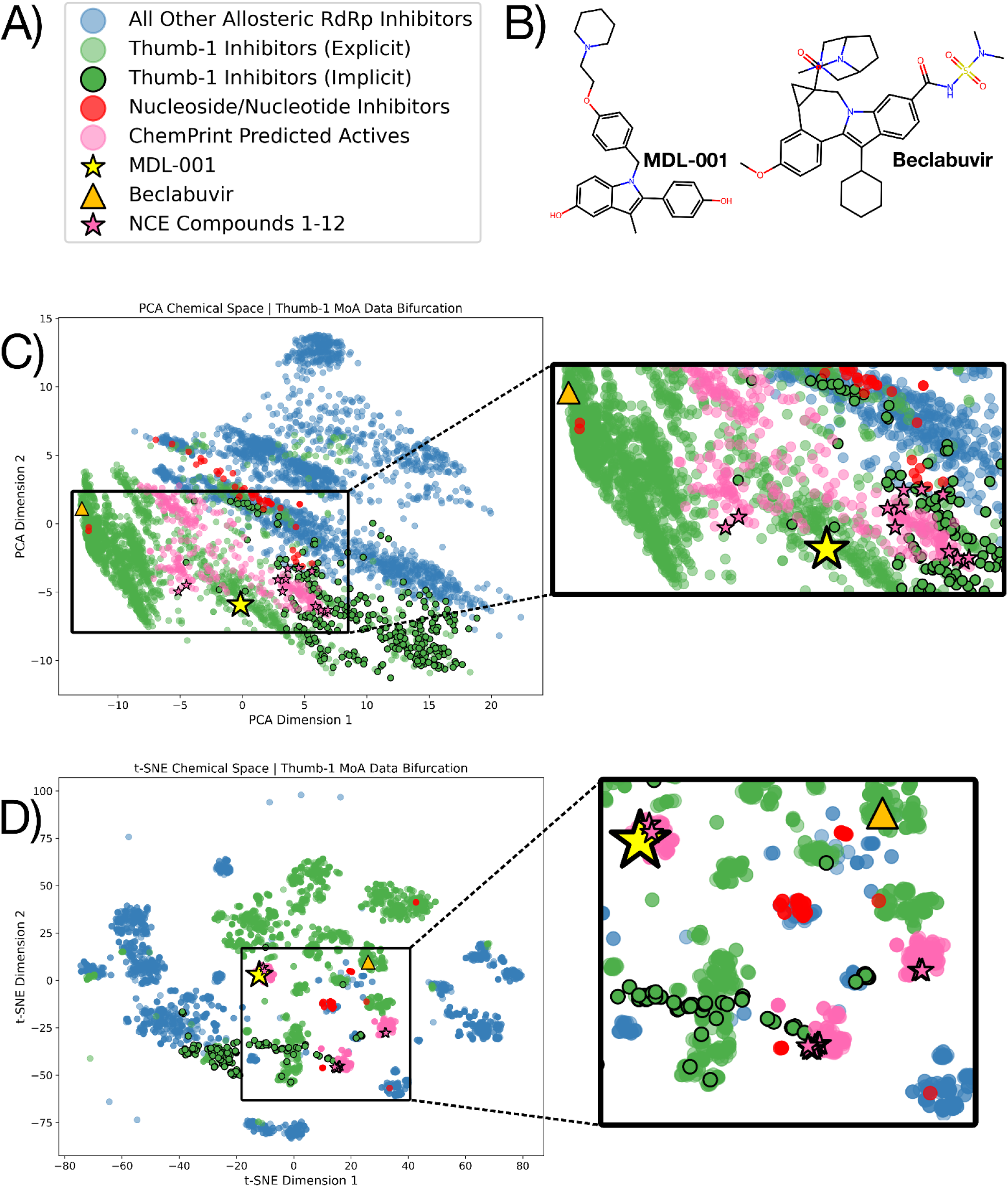
**A)** Label names, color code, and symbols assigned to the data visualized with PCA and t-SNE. Dataset composition and label logic is explained in materials and methods. **B)** Molecular structures of MDL-001 and Beclabuvir; **C)** PCA where the first two principal components explain 24% of the total variance in the data and **D)** t-SNE of the chemical space for broad-spectrum antiviral ligands. The rectangular inset in panels **C** and **D**, show the chemical space covered by the 12 NCEs, MDL-001, Beclabuvir in comparison to the curated datasets.

PCA (**Figure 2C**) revealed a significant pattern of chemical distribution associated with documented MoAs. MDL-001 and the ChemPrint-predicted compounds, including the 12 NCEs, occupy a chemical space between ‘explicit’ inhibitors, known for targeting HCV’s RdRp Thumb-1 pocket, and ‘implicit’ inhibitors, which target broader viral families without direct Thumb-1 evidence but align with its pharmacophoric profile or core scaffolds. Beclabuvir, in contrast, occupies a distinct corner of chemical space only occupied by ‘explicit’ HCV Thumb-1 inhibitors.

t-SNE (**Figure 2D**), which better preserves local relationships, revealed clustering consistent with labeled MoAs. Three distinct local clusters containing the ChemPrint-predicted compounds, including the 12 NCEs, occupy the space near ‘explicit’ and ‘implicit’ inhibitors. One of these three clusters includes MDL-001, while none include Beclabuvir.

Tanimoto coefficients provided quantitative evidence of chemical similarity between the 12 NCEs vs MDL-001 and Beclabuvir (**Figure 3**). Values range from 0 to 1, with a Tanimoto coefficient of ≥ 0.5 (using ECFP4) indicating chemical similarity between two small molecules. The 12 NCEs received Tanimoto coefficients ranging from 0.28–0.47 compared to MDL-001, with an average similarity of 0.38, indicating chemical dissimilarity. When compared to Beclabuvir, the Tanimoto coefficients were even lower, ranging from 0.10–0.20, with an average similarity of 0.13. All NCE compounds reported scores ≤0.47 compared to MDL-001 and ≤0.20 compared to Beclabuvir, demonstrating significant chemical novelty.

**Figure 3.**
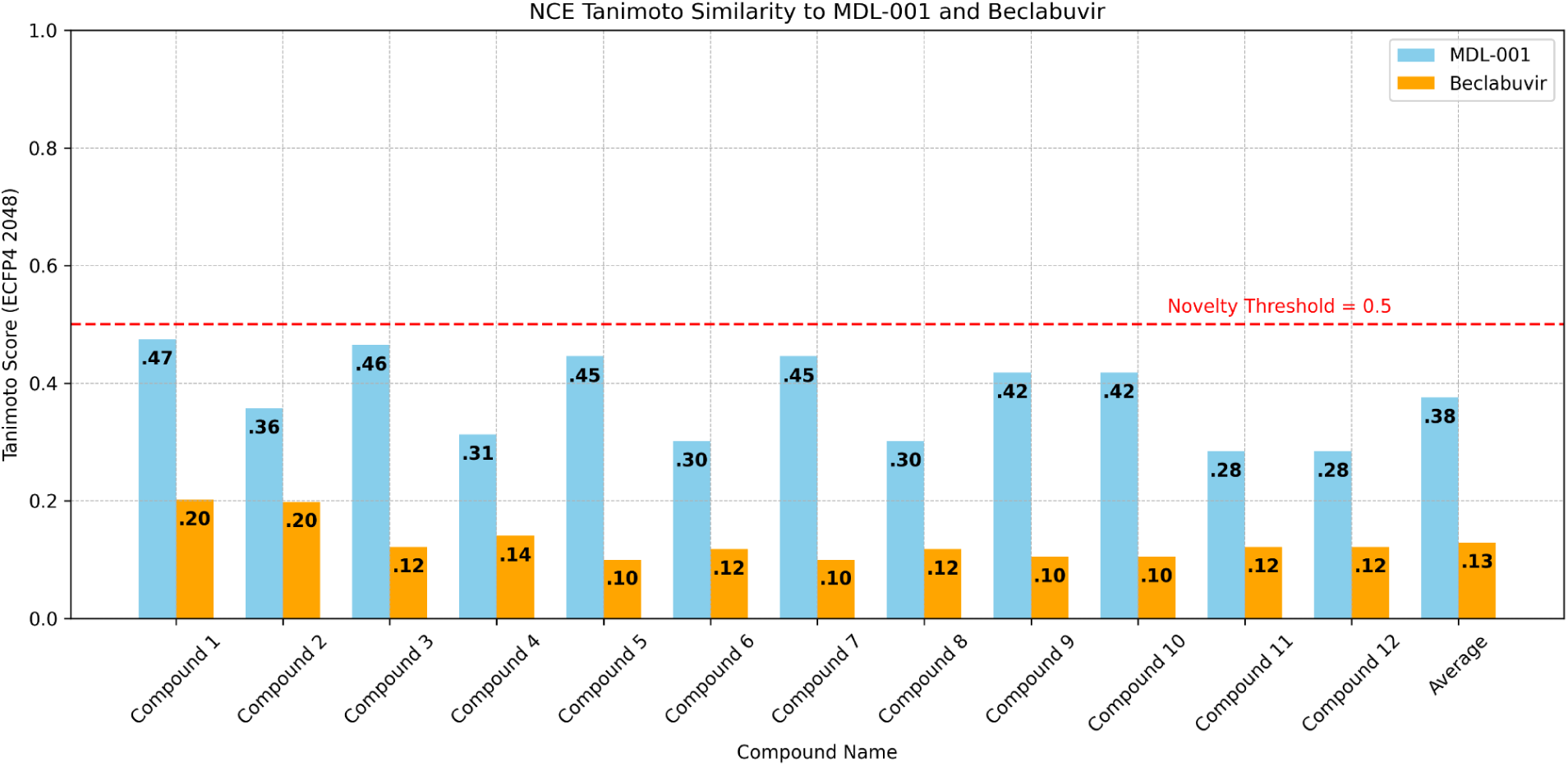
Tanimoto coefficients. The grouped bar plot in this figure shows the Tanimoto coefficients calculated for the 12 NCEs, when compared to MDL-001 (light blue bars) and Beclanbuvir (orange bars). Small molecules were embedded in the ECFP4 format for Tanimoto coefficient calculation. The dashed, horizontal line (colored red) indicates the chemical novelty threshold (Tanimoto coefficient ≥ .5).

### Compound synthesis

The 12 NCEs selected for experimental validation were successfully synthesized at Piramal Pharma Solutions, with all compounds achieving a purity greater than 95%. This ensured their suitability for downstream validation of bioactivity.

### *In vitro* broad-spectrum antiviral potency

To validate NCE antiviral activity experimentally, we used pseudovirus assays specific to HCV 1b and human Coronavirus 229E (CoV229E) (**Figure 4** and **Table 2**). In these assays, all 12 NCEs showed antiviral activity against either HCV or CoV229E, with EC_50_ values ranging from 0.29-10 μM, though CoV229E was more sensitive than HCV. Specifically, 11 of 12 NCEs inhibited the Cov229E pseudogene. On the other hand, 9 of 12 NCEs inhibited the HCV pseudovirus.

**Figure 4.**
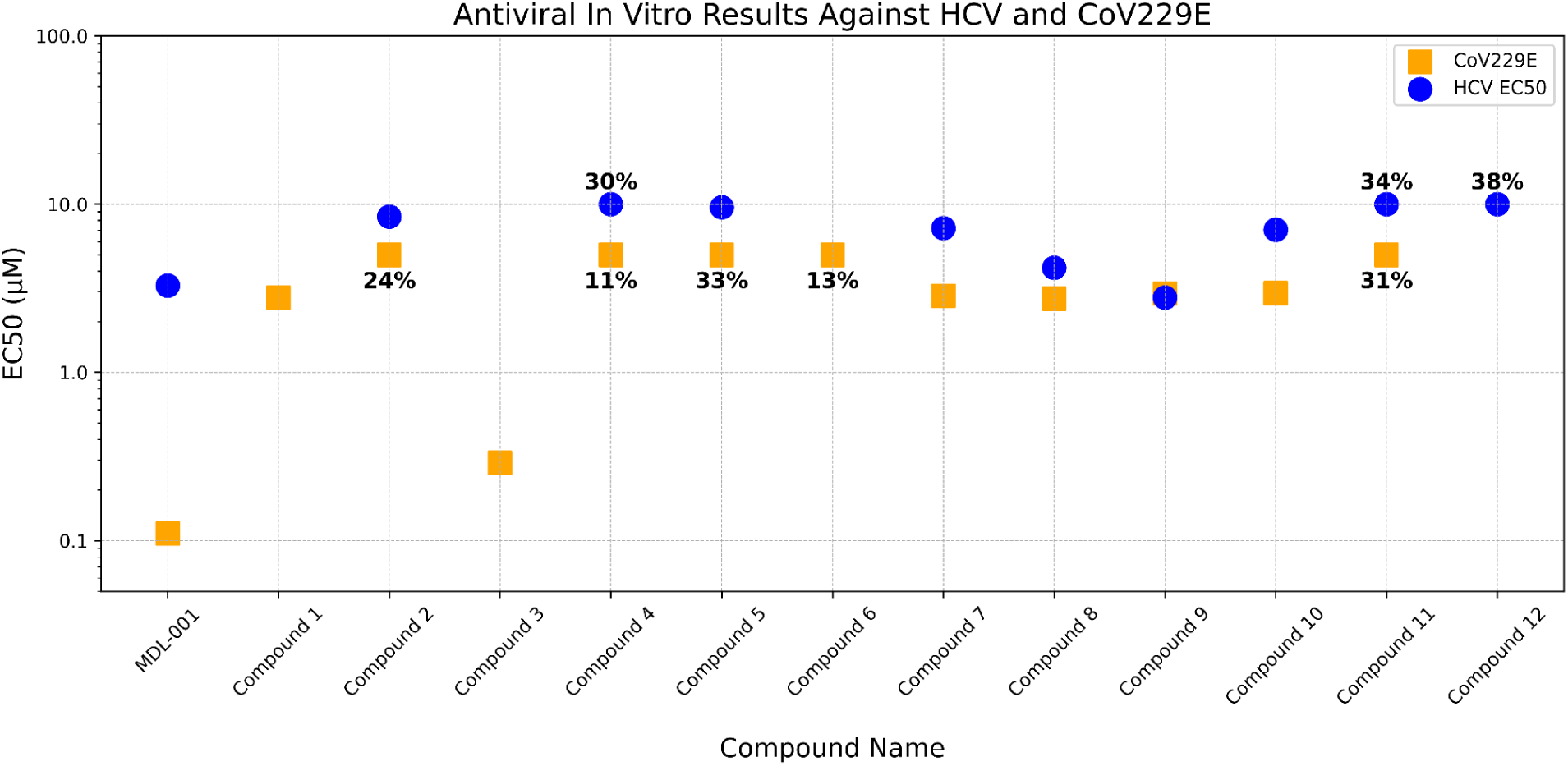
*In vitro* antiviral assays. Shown in this figure are the EC_50_ values obtained for the 12 NCEs and MDL-001, when tested against pseudovirus inhibitory assays specific to HCV 1b (blue circles) and CoV229E (orange squares). EC_50_ values are plotted in μM units along the y-axis.

**Table 2.** *In vitro* results of twelve compounds against HCV 1b and CoV229E.

| Compound ID | HCV 1b - EC<sub>50</sub> (μM) | HCV 1b - TC<sub>50</sub> (μM) | CoV229E - EC<sub>50</sub> (μM) | CoV229E - TC<sub>50</sub> (μM) |
| --- | --- | --- | --- | --- |
| MDL-001 | 3.28 | >10 | 0.11 | >1 |
| Compound 1 | - | >10 | 2.78 | >5 |
| Compound 2 | 8.43 | >10 | 24% @ 5 μM | 2.47 |
| Compound 3 | - | >10 | 0.29 | 1.53 |
| Compound 4 | 30% @ 10 μM | >10 | 11% @ 5 μM | >5 |
| Compound 5 | 9.59 | >10 | 33% @ 5 μM | >5 |
| Compound 6 | - | >10 | 13% @ 5 μM | >5 |
| Compound 7 | 7.2 | >10 | 2.85 | >5 |
| Compound 8 | 4.19 | 5.82 | 2.74 | >5 |
| Compound 9 | 2.78 | >10 | 2.94 | >5 |
| Compound 10 | 7.05 | >10 | 2.96 | >5 |
| Compound 11 | 34% @ 10 μM | >10 | 31% @ 5 μM | >5 |
| Compound 12 | 38% @ 10 μM | >10 | - | >5 |

### Estrogen receptor-dependent cell growth inhibition

We evaluated the 12 NCEs for selective estrogen regulation modulation activity by using a growth inhibition assay. The assay is performed on MCF7 cells, which are derived from a hormone-dependent breast cancer that stops growing upon estrogen receptor inhibition. Our results revealed that, in contrast to MDL-001, the 12 NCEs lacked MCF7 growth inhibition properties (**Figure 5** and **Table 3**).

**Figure 5.**
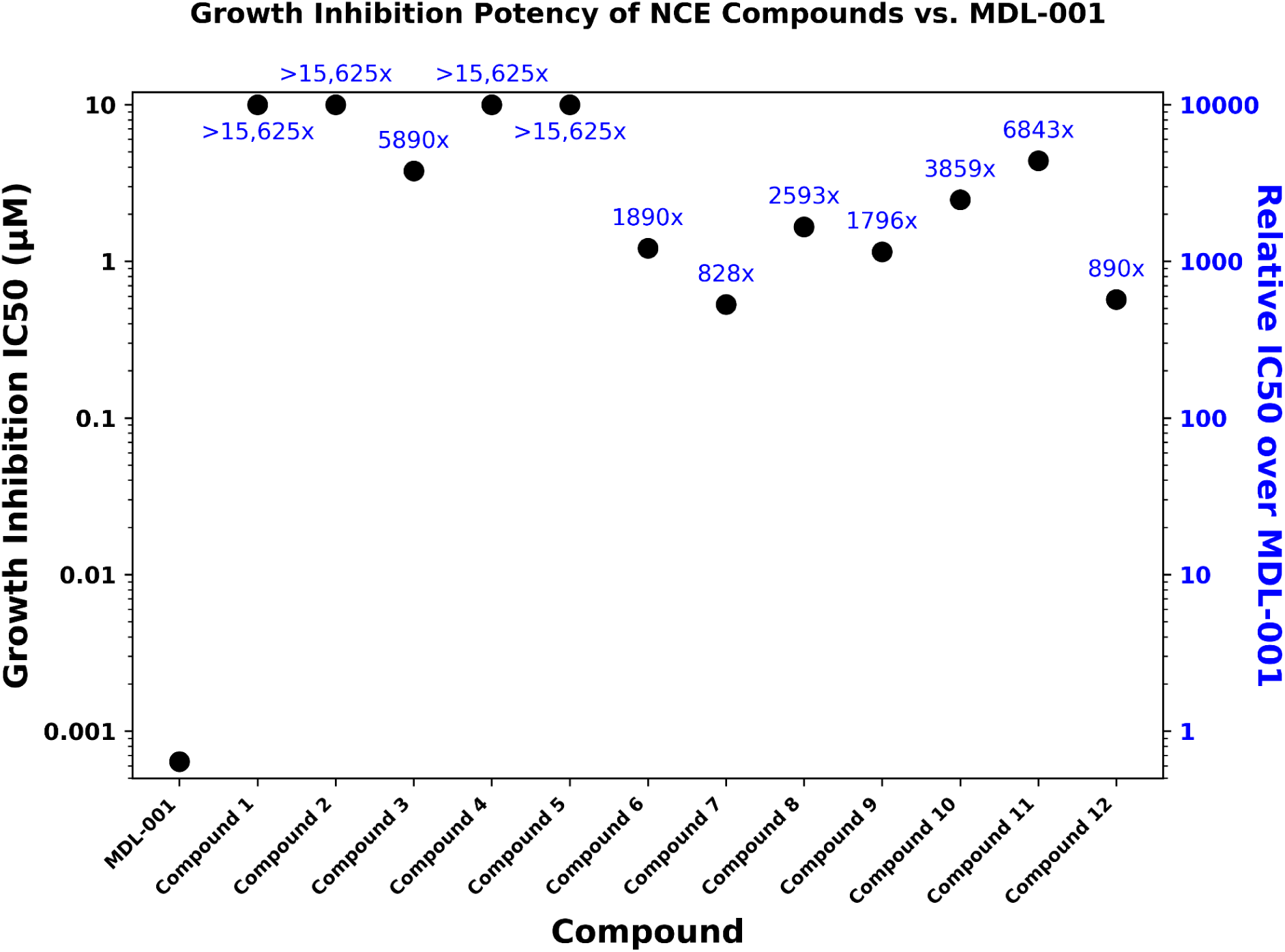
MCF7 growth inhibition assays. The plot in this figure shows the half-maximal inhibitory concentration (IC_50_) values for the 12 NCEs and MDL-001, when added to MCF7 cells. IC_50_ values are plotted along the left y-axis. The relative NCE IC_50_ with respect to MDL-001’s is plotted along the right y-axis.

**Table 3.**
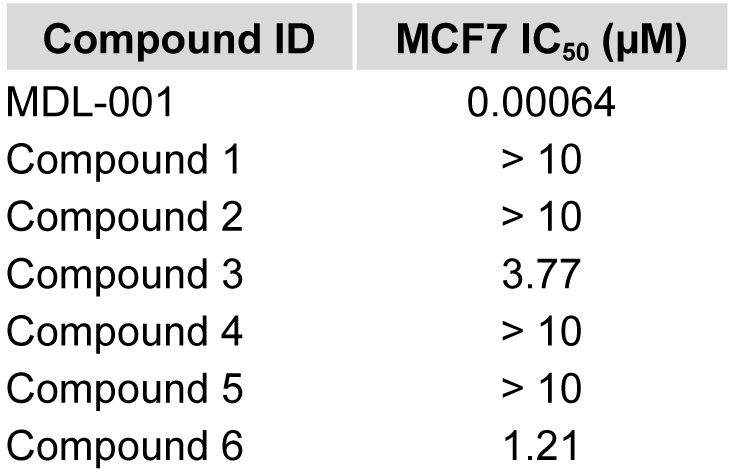
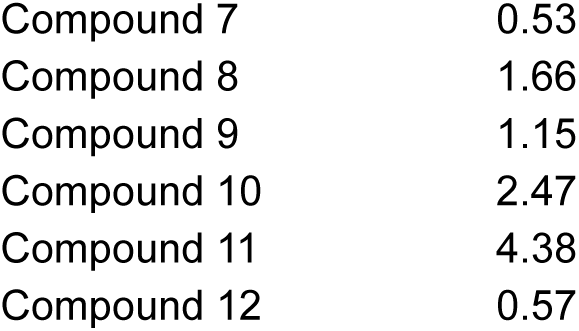
*In vitro* results of twelve compounds in MCF7 growth inhibition assay.

| Compound ID | MCF7 IC<sub>50</sub> (µM) |
| --- | --- |
| MDL-001 | 0.00064 |
| Compound 1 | > 10 |
| Compound 2 | > 10 |
| Compound 3 | 3.77 |
| Compound 4 | > 10 |
| Compound 5 | > 10 |
| Compound 6 | 1.21 |
| Compound 7 | 0.53 |
| Compound 8 | 1.66 |
| Compound 9 | 1.15 |
| Compound 10 | 2.47 |
| Compound 11 | 4.38 |
| Compound 12 | 0.57 |

## Discussion

This study highlights the productivity of GALILEO as a one-shot, multimodal drug discovery pipeline. Building on the established success of MDL-001, a potential first-in-class broad-spectrum antiviral therapeutic with demonstrated efficacy and safety, GALILEO developed a next-generation library of antiviral NCEs with enhanced target specificity. Using GALILEO’s generative tools, we explored chemical space containing trillions of molecules derived from MDL-001’s pharmacophoric scaffold. Through a multimodal approach that simultaneously optimized antiviral activity and target specificity, ChemPrint’s one-shot predictions identified 12 NCEs with broad-spectrum antiviral activity and significantly enhanced target specificity relative to MDL-001. The identified NCEs showed *in vitro* activity against Hepatitis C Virus (HCV) and/or human Coronavirus 229E, achieving a 100% hit rate. *In vitro* studies confirmed that the NCEs exhibited specificity improvements of 800-fold to greater than 15,000-fold relative to MDL-001’s original target. These results highlight GALILEO’s capacity to produce highly optimized drug candidates with exceptional efficiency.

The 12 NCEs share four attributes: a) high Thumb-1 binding probability; b) low estrogen receptor binding probability; c) broad-spectrum antiviral activity; and d) negligible, if any, inhibition of estrogen receptor-mediated growth. Tanimoto similarity analysis demonstrated the chemical novelty of the NCE library, with low similarity to MDL-001 (average Tanimoto coefficient: 0.38) and Beclabuvir (average Tanimoto coefficient: 0.13), as well as to a curated library of known antiviral drugs. Chemical space mapping further revealed that these NCEs occupy a distinct region, highlighting their uniqueness within the context of existing antiviral therapeutics and GALILEO’s one-shot ability to identify biologically relevant compounds in unexplored areas of chemical space.

By integrating generative and multimodal methodologies, GALILEO addresses critical challenges in antiviral drug discovery, providing an efficient pathway to identify and optimize novel, broad-spectrum candidates. The 12 NCEs identified hold great therapeutic value, which must be investigated in detail. We expect favorable therapeutic properties (oral availability, desirable pharmacokinetics, and clinical safety) for these 12 NCEs, because their chemical structure is based on MDL-001’s pharmacophoric scaffold. Future studies will assess their pharmacological properties in preclinical settings and their broad-spectrum activity in a large battery of viruses. If needed, lead optimization will be conducted using GALILEO to further refine their efficacy and selectivity. While demonstrated here in the development of broad-spectrum antiviral drugs, GALILEO’s design is versatile and can support AI-driven drug discovery across diverse therapeutic areas.

## Methods

### Broad-spectrum training data

The ChemPrint dataset, which we described in detail previously, comprises a set of small molecules with well-documented antiviral activity.³ This dataset included small molecules active against four viral families: *Coronaviridae*, *Picornaviridae*, *Flaviviridae*, and *Togaviridae*. Antiviral compounds were labeled as “explicit” if their Thumb-1 site inhibition in HCV’s RdRp was directly identified in the literature. Compounds were labeled as "implicit" if they targeted the RdRp of any ssRNA virus without direct evidence of Thumb-1 binding but matched the Thumb-1 pharmacophoric profile or contained core scaffolds consistent with Thumb-1 inhibitors. Both explicit and implicit inhibitors were included in the broad-spectrum training dataset.

### Broad-spectrum training data processing

Small molecules were removed from the dataset if their size fell outside a 180-800 Da size range, if they lacked K_i_, K_d_, IC_50_ or EC_50_ values, or if they contained a PAINS (Pan-Assay Interference Compounds) substructure, a category of molecules documented for their non-specific activity.^20^ Small molecules with an ‘indol_3yl_alk(461)’ substructure were kept because this moiety is commonly found in antiviral drugs. A binary classification dataset was then created by defining a line of best fit across viral datasets to establish a relationship between negative log bioactivity measures. Missing values were imputed using this relationship, and an orthogonal line to the best fit determined the activity cutoff. Compounds were classified as active if they inhibited two or more viruses or, in cases of limited testing, a single virus. This process resulted in 1,282 compounds, with 602 classified as active and 680 as inactive.

### Estrogen receptor training data

We used ChemPrint to identify and remove NCEs predicted to interact with the human estrogen receptor. The dataset comprised small molecule ligands known to bind ERα (Uniprot ID: P03372) and ERβ (Uniprot ID: Q92731). Bioactivity data for these ligands were retrieved from PubChem, BindingDB, and ChEMBL, including compounds classified as agonists, antagonists, or selective estrogen receptor modulators (SERMs). Using data from all these categories was appropriate because they share the common feature of binding to the estrogen receptor, enabling comprehensive identification of compounds with potential receptor interaction.

### Estrogen receptor training data processing

As was done with the Thumb-1 training dataset, small molecules were removed if their size fell outside the 180–800 Da range, if they lacked K_i_, K_d_, IC_50_ or EC_50_ values, or if they contained a PAINS (Pan-Assay Interference Compounds) substructure, excluding ‘indol_3yl_alk(461)’.

Duplicates were resolved by retaining the most potent readout, irrespective of the type of bioactivity measurement or ERα/ERβ interaction. An activity cutoff of ∼75 nM was applied to classify compounds as active or inactive, creating a balanced dataset. This threshold was selected to identify compounds with lower potency relative to MDL-001, which has an IC50 of 14 nM against ERα. The final dataset included 4,208 compounds, with 2,103 classified as active and 2,105 as inactive.

### ChemPrint model architecture

ChemPrint architecture, previously detailed by us,^5^ leverages Mol-GDL to learn the adaptive embeddings derived from its training data to make predictions.^21^ The input data takes the form of a geometric graph, a molecular representation that encapsulates the structural information of each datapoint, to which one-hot encoded features are passed in. ChemPrint architecture encompasses an end-to-end Graph Convolutional Network (GCN) with a Multilayer Perceptron (MLP) module to facilitate positive and negative classification. Select normalization, activation, pooling, and dropout layers were used.

### Model training for Thumb-1 interactors

Datasets were split 50/50 into training and validation sets, with the t-SNE split methodology, we reported previously.^5^ Receiver operating characteristic curves (AUROC) to assess the model’s performance during hyperparameter optimization (**Figure S1**). For large-scale inference, a single broad-spectrum model optimized with the full dataset was deployed.

### Model training for estrogen receptor interactors

Datasets were split and model performance were achieved as described in the above subsection for the model trained to predict Thumb-1 interactors (**Figure S2**). In this case inference was accomplished using an ensemble of 10 estrogen receptor ChemPrint models, retrained on the full dataset.

### Broad-spectrum combinatorial space

The combinatorial chemical space was generated from MDL-001’s pharmacophoric scaffold and amounted to 52 trillion unique small molecules. Each of these small molecules was decorated with up to five R group substituents, designed to engage in hydrogen bond interactions with the Thumb-1 pocket. The scaffolds and substituent positions were informed by training data targeting the Thumb-1 site and synthesizability.

### Broad-spectrum inference space

The broad-spectrum inference library was generated by reducing the combinatorial space to approximately one billion compounds. This was achieved by removing stereospecificity from all substituents (R1–R5) and simplifying substituent structures based on synthetic accessibility. Adjustments included the removal of certain alkyl groups and reducing the complexity of R group combinations. These steps allowed for a more manageable and focused prediction library, while retaining chemistry representative of the entire combinatorial space.

### Model inference for Thumb-1 interactors

The broad-spectrum Thumb-1 ChemPrint model was deployed for brute force inference on approximately one billion compounds. Data featurization and inference required ∼2,000 computational hours, utilizing multiprocessing on a system with 48 CPU cores. By leveraging 100 CPUs, each with 48 cores operating in parallel, we were able to complete the screening of one billion compounds in under 24 hours.

### Model inference for estrogen receptor interactors

An ensemble of 10 estrogen receptor ChemPrint models was applied to the refined list of 500 NCEs filtered by the broad-spectrum Thumb-1 ChemPrint model. This inference step was performed to identify and exclude compounds predicted to interact with the estrogen receptor.

### Chemical space data

The datasets used for chemical space analysis comprised small molecules with well-documented antiviral activity, as described previously.³ These included compounds active against the viral families Coronaviridae, Picornaviridae, Flaviviridae, and Togaviridae. Compounds were classified as ‘explicit’ inhibitors if their activity was directly linked to the Thumb-1 pocket in HCV’s RdRp. Compounds were classified as ‘implicit’ inhibitors if they targeted viruses without direct evidence of Thumb-1 binding but matched the Thumb-1 pharmacophoric profile or contained core scaffolds consistent with Thumb-1 inhibitors. Additional data included nucleoside/nucleotide inhibitors (active site inhibitors), other allosteric RdRp inhibitors (not targeting Thumb-1), MDL-001, Beclabuvir, a subset of ChemPrint-predicted broad-spectrum Thumb-1 active compounds, and the 12 NCEs.

### Chemical space mapping with PCA and t-SNE

Principal component analysis (PCA) and t-distributed stochastic neighbor embedding (t-SNE) were used to visualize chemical space relationships. Both methods used chemical structures as input features, represented as ECFP4 fingerprints. No mechanism of action (MoA) labels were used as input features for the analysis. PCA, a linear dimensionality reduction technique, captured global relationships in the dataset, with the distance between points reflecting structural similarity. In contrast, t-SNE, a non-linear technique, better preserved local relationships, providing insight into clustering within the chemical space. The results of PCA and t-SNE were visualized in two-dimensional plots to compare the relative positioning of MDL-001, Beclabuvir, the 12 NCEs, ChemPrint-predicted compounds, and curated antiviral datasets.

### Tanimoto Similarity Analysis

Tanimoto coefficients were calculated to quantify chemical similarity between compounds. ECFP4 fingerprints were used to compute pairwise similarity scores, ranging from 0 to 1. Two compounds were considered chemically similar if the Tanimoto coefficient was ≥ 0.5, a threshold commonly used in the industry to denote significant similarity. Comparisons were performed between the 12 NCEs, MDL-001, and Beclabuvir to assess their relative novelty and diversity within the chemical space.

### Compound synthesis

The 12 NCEs were synthesized at Piramal Pharma Solutions. Purity was confirmed to be greater than 95% for all compounds, verified through liquid chromatography–mass spectrometry (LC–MS), high-performance liquid chromatography (HPLC), and proton nuclear magnetic resonance (¹H-NMR) spectroscopy. These methods ensured the chemical integrity and suitability of the synthesized compounds for subsequent testing and analysis.

### *In vitro* antiviral assays Cell culture

The Huh-lucineo-ET reporter cell line was obtained from Dr. Ralf Bartenschlager (Department of Molecular Virology, Hygiene Institute, University of Heidelberg, Germany) by ImQuest BioSciences through a specific licensing agreement. This cell line harbors the persistently replicating 1389/NS3-37LucUbiNeo-ET replicon of HCV genotype 1b containing the firefly luciferase gene-ubiquitin-neomycin phosphotransferase fusion protein and EMCV IRES driven N53-5B HCV coding sequences containing the ET tissue culture adaptive mutations (E1202G, T12081, and K1846T). For the first two passages after thawing, the cells were cultured in DMEM without phenol red (Gibco, catalog no. 31053028) supplemented with fetal bovine serum (FBS) (Gibco, catalog no. 10082147), 2 mM GlutaMAX Supplement (Gibco, catalog no. 35050061), 100 U/mL penicillin and 100 µg/mL streptomycin (Corning, catalog no. 30-002-CI), and 1X nonessential amino acids (Gibco, catalog no. 11140050) plus 1 mg/mL G418 (Gibco, catalog no. 10131035). The cells were then further cultured in the complete medium described above plus 500 µg/mL G418. One day prior to test article addition, the cells were seeded at a density of 5 x 103 cells per well in 100 µL complete medium plus 500 µg/mL G418 in either standard clear or white-walled/clear-bottom 96-well tissue culture plates for colorimetric or chemiluminescent endpoint assays, respectively. The cells were incubated at 37°C/5% CO2 for 24 hours.

### Huh-lucineo-ET treatment conditions

Twenty-four hours after seeding the cells, the cell culture supernatant was removed from each well and replaced with 175 µL fresh complete medium minus G418. Test compounds were evaluated at a high-test concentration of 10 µM and five serial half-logarithmic dilutions in three biological replicates. In three biological replicates, Sofosbuvir was evaluated in parallel at a high-test concentration of 1 µM and five serial half-logarithmic dilutions. Compounds were first serially diluted in DMSO to 400X of each final in-well concentration. An intermediate 8X solution was prepared in a complete medium and then added to the cells in a volume of 25 µL per well. A complete medium containing an equal percentage of DMSO (v/v) was added to vehicle-treated control cells. The final DMSO concentration was 0.25% (v/v) for all samples. The cells were incubated for an additional 72 hours at 37°C/5% CO2, then HCV replication was measured by luciferase activity and cell viability was assessed in parallel using an XTT colorimetric assay.

### Measurement of HCV replication

HCV replication from the replicon assay system was measured following 72 hours of incubation using the britelite plus Reporter Gene Assay according to the manufacturer’s protocol (Revvity, catalog no. 6066761). In brief, one vial of britelite plus substrate was solubilized in 10 mL of britelite reconstitution buffer, mixed gently by inversion, and incubated for five minutes at room temperature. One hundred microliters (100 µL) of cell culture supernatant were removed from each well, followed by adding 100 µL britelite plus reagent and thorough mixing by pipetting up and down. The plates were sealed with adhesive film and incubated at room temperature for 10 minutes. Chemiluminescence was measured as relative light units on a FlexStation 3 microplate reader (Molecular Devices). The raw data were imported into Microsoft Excel for further analysis. Data are presented as relative light units (RLU) or percent relative to vehicle-treated, virus-only control (mean +1- standard deviation, n=3 biological replicates from one independent experiment).

### XTT cell viability assay

Huh-luc/neo-ET cell viability was measured at 72 hours after treatment using the tetrazolium dye XTT (2,3-bis(2-methoxy-4-nitro-5-sulfophenyI)-5[(phenylamino)carbonyl]-2H- tetrazolium hydroxide). Mitochondrial enzymes of metabolically active cells metabolize XTT-tetrazolium to a soluble formazan product. XTT solution was prepared fresh at 1 mg/mL in DMEM. Phenazine methosulfate (PMS) solution was prepared at 0.15 mg/mL in PBS and stored in the dark at −20°C. An XTT/PMS stock was prepared immediately before use by adding 40 µL of PMS per ml of XTT solution. Fifty microliters (50 µL) XTT/PMS were added per well, and the cells were incubated for 4 hours at 37°C. The cell culture plates were then sealed with adhesive plate sealers and shaken gently or inverted several times to mix the soluble formazan product. The formazan product was measured spectrophotometrically by absorbance at 450/650 nm on a Vmax plate reader (Molecular Devices). The raw data in Softmax Pro 5.4.2 software was imported into Microsoft Excel for further analysis. Data are presented as absorbance values (450/650 nm) or the percentage of viable cells relative to vehicle-treated control (mean +1- standard deviation, n=3 biological replicates from one independent experiment).

### *In vitro* activity and anti-Coronavirus 229E cytoprotection assay

#### Cell preparation

MRC-5 cells obtained from ATCC (CCL-171) were passaged in DMEM supplemented with 10% FBS, 2 mM L-glutamine, 100 U/mL penicillin, 100 µg/mL streptomycin, 1 mM sodium pyruvate, and 0.1 mM NEAA in T-75 flasks prior to use in the antiviral assay. On the day preceding the assay, the cells were split 1:2 to ensure they were in an exponential growth phase at the time of infection. Total cell and viability quantification was performed using a hemocytometer and Trypan Blue dye exclusion. Cell viability was greater than 95% for the cells to be utilized in the assay. The cells were resuspended at 3 x 103 cells per well in a tissue culture medium and added to flat bottom microtiter plates in a volume of 100 µL. The plates were incubated at 37°C/5% CO2 overnight to allow for cell adherence.

### Virus preparation

Coronavirus229E (CoV229E) was obtained from ATCC (VR-740) and grown in MRC-5 cells to produce a stock virus pool. A pretittered aliquot of the virus was removed from the freezer (−80°C) and allowed to thaw slowly to room temperature in a biological safety cabinet. The virus was resuspended and diluted into assay medium (DMEM supplemented with 2% heat-inactivated FBS, 2 mM L-glutamine, 100 U/mL penicillin, 100 µg/mL streptomycin) such that the amount of virus added to each well in a volume of 100 µL was the amount determined to yield 85 to 95% cell killing at 6 days post-infection.

### Plate format

Each plate contains cell control wells (cells only), virus control wells (cells plus virus), triplicate drug toxicity wells per compound (cells plus drug only), and triplicate experimental wells (drug plus cells plus virus).

### Efficacy and toxicity XTT

Following incubation at 37°C in a 5% CO2 incubator for six days, the test plates were stained with the tetrazolium dye XTT (2,3-bis(2-methoxy-4-nitro-5-sulfophenyl)-5- [(phenylamino)carbonyl]-2H-tetrazolium hydroxide). The mitochondria enzymes of metabolically active cells metabolized XTT-tetrazolium to a soluble formazan product, allowing rapid quantitative analysis of the inhibition of virus-induced cell killing by antiviral test substances. XTT solution was prepared daily as a 1 mg/mL stock in RPMI1640. Phenazine methosulfate (PMS) solution was prepared at 0.15 mg/mL in PBS and stored in the dark at −20°C. XTT/PMS stock was prepared immediately before use by adding 40 µL of PMS per ml of XTT solution. Fifty microliters of XTT/PMS was added to each well of the plate, and the plate was reincubated for 4 hours at 37°C. Plates were sealed with adhesive plate sealers and shaken gently or inverted several times to mix the soluble formazan product. The plate was read spectrophotometrically at 450/650 nm with a Molecular Devices V_max_ plate reader.

### MCF7 growth inhibition assays

#### Assay conditions

The growth inhibition properties of inhibitors on MCF7 wild-type cell lines were evaluated using the Cell Titre Glo 2.0 cell viability reagent. The assay utilized 384-well black clear bottom plates with 20 µl of MCF7 wild-type cell suspension mixed with reference/test compounds and 20 µl of Cell Titre Glo 2.0 reagent. Cells were seeded at a density of 1000 cells/well in DMEM High Glucose culture media + 10% Fetal Bovine Serum (FBS). Staurosporine or Camptothecin served as a positive control for confirming cytotoxicity at a concentration of 30 µM, and vehicle control of 1% DMSO was used to standardize conditions across all test points. The assay and culture media both contained 1% DMSO. Assays were conducted in duplicate for each concentration of the test compounds. The incubation period for the assay was set at 120 hours, after which growth inhibition was assessed. Using a multi-mode reader, the assay readout was based on luminescence measured in Relative Light Units (RLUs). The viability luminescence signal was monitored continually from the same wells over an extended period, allowing for detailed analysis of cell viability. Measurements were taken using an integration time ranging from 0.25 to 1 second across plates designated for various time points.

#### Assay protocol

Cells were maintained and seeded in 384 healthy plates, maintaining 1000 cells/well. Seeded plates were incubated at 37°C, 5% CO2 for 16-18 hours to ensure optimal cell adherence and growth. Post-incubation, reference, and test compounds were added carefully to the wells for further processing. Plates were incubated again at 37°C, 5% CO2 for 120 hours (5 days) to allow compound effects. This was followed by the addition of CTG reagent as per the kit’s instruction protocol for 30 minutes and RLU data to be recorded. Post plate read data to be analyzed using GraphPad Prism for software IC_50_ value generation.

