## Supplemental Figures S1 and S2 for "GALILEO Generatively Expands Chemical Space and Achieves One-Shot Identification of a Library of Novel, Specific, Next Generation Broad-Spectrum Antiviral Compounds at High Hit Rates"

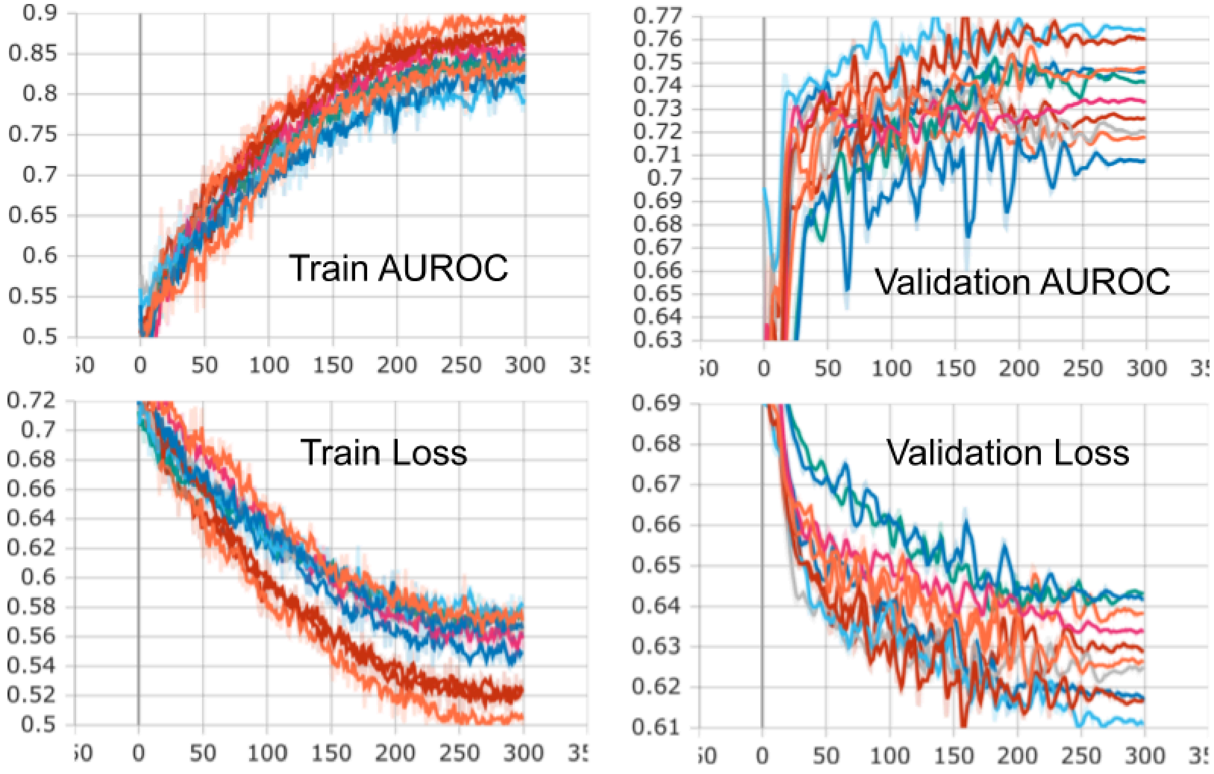


**Figure S1.** AUROC and binary cross-entropy loss for t-SNE split training and validation sets (broad-spectrum model).

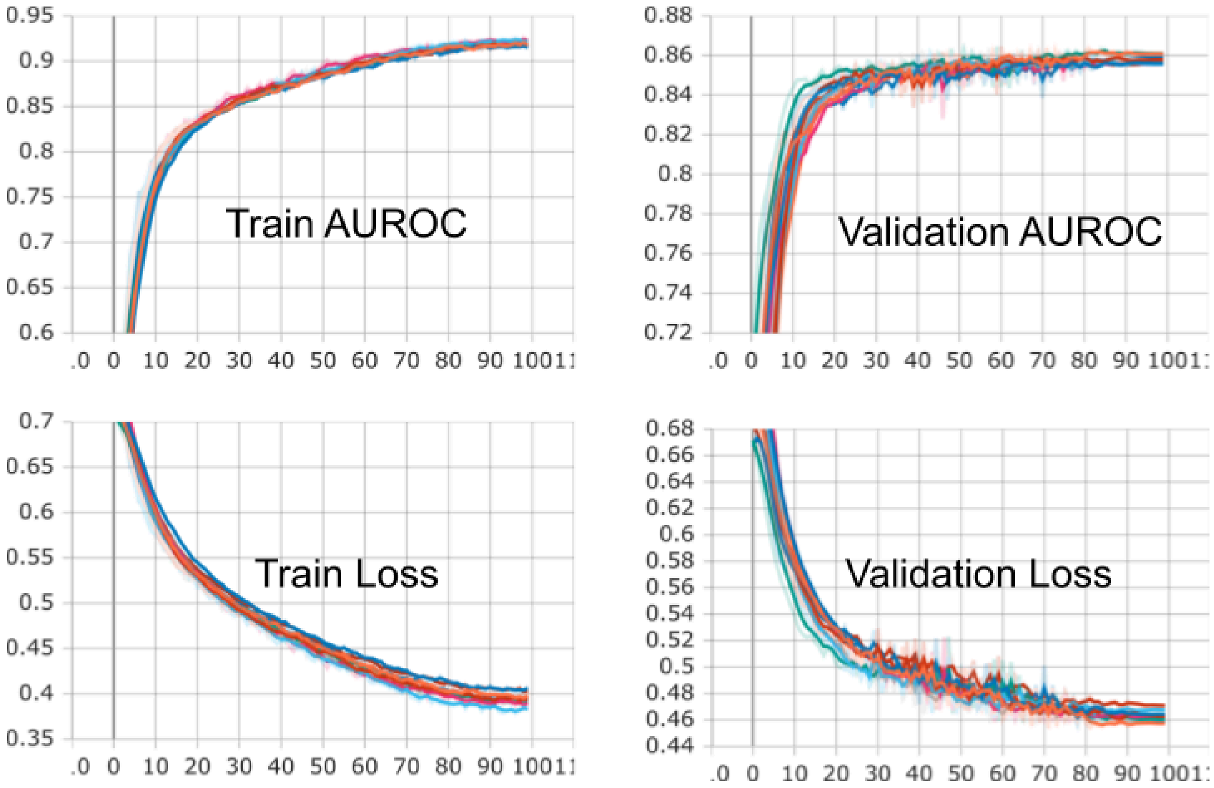


**Figure S2.** AUROC and binary cross-entropy loss for t-SNE split training and validation sets (ER binding model).
